# Metagenomic exploration of the bacteriome reveals natural *Wolbachia* infections in yellow fever mosquito *Aedes aegypti* and Asian tiger mosquito *Aedes albopictus*

**DOI:** 10.1101/2025.03.06.641799

**Authors:** Gul E Nayab, Rafi ur Rahman, Fazal Hanan, Inamullah Khan, Muhammad Fahim

## Abstract

**Background:** Dengue and associated complications are spreading to non-endemic regions of Pakistan. Vector control, the foremost and widely adopted strategy for managing dengue has been implemented through various measures in Pakistan. Biological control through the use of *Wolbachia*, a bacterium naturally present in various insect genera, including *Aedes*, has demonstrated promising results globally. In this study we collected *Aedes* species and investigated their microbiomes with a particular focus on identifying the endosymbiont *Wolbachia*.

**Methods:** Mosquitoes were collected via Gravitraps in the Peshawar region of northwest Pakistan. The identity of the mosquitoes was initially confirmed through morphological characters followed by molecular identification using species-specific Cytochrome oxidase I (COI) primers. The DNA from female *Ae. aegypti* and *Ae. albopictus* was further subjected to 16S rRNA sequencing. The hypervariable regions V3/V4 of 16S rRNA were used for sequencing using the paired-end Illumina MiSeq platform.

**Results:** The phylogenetic analysis of the COI gene in our samples demonstrated similarity to *Aedes* species previously documented in Pakistan. In comparative analysis of their microbiomes, *Ae. albopictus* was found to harbor 921 bacterial species, while *Ae. aegypti* only had 239 species. The metagenomic analysis revealed *Wolbachia pipientis* infection in *Ae. aegypti* while co-infection of *Wolbachia pipientis* and *Wolbachia bourtzisii* was detected in *Ae. albopictus* microbiota.

**Conclusion:** Both *Ae. aegypti* and *Ae. albopictus* are present in Peshawar region of Khyber Pakhtoonkhwa province of Pakistan. Comparative analysis of the bacteriome showed higher bacterial diversity for *Ae. albopictus* as compared to *Ae. aegypti.* The investigation revealed *Wolbachia* infection in both *Aedes* mosquitoes species.

## Introduction

Vector-borne diseases (VBDs) impose a significant burden of morbidity and mortality worldwide, placing considerable strain on public health systems [1, 2]. Dengue, a crucial VBD, is caused by a *Flavivirus* and transmitted to humans by the bite of an infected female *Aedes* mosquito [3, 4]. Every year, approximately 4 billion people, nearly half of the global population, are at risk of contracting dengue infection, predominantly in tropical and subtropical regions [5, 6]. In Pakistan, dengue first emerged in 1994 [7]. Since 2005, dengue has become an endemic disease in the country, posing a persistent threat to millions of lives annually [8, 9]. Records of dengue outbreaks have been reported consequently in 2011, 2013 and 2015 from Punjab, Sindh and Khyber Pakhtunkhwa provinces respectively [10, 11]. During the 2011 dengue outbreak, around 20,000 infections and almost 300 deaths were reported in Lahore city alone [12]. The Northern Swat district reported eight thousand cases and 57 deaths in the outbreak of 2013 [10]. Every year, post-monsoon (September-October) sees a surge in nationwide dengue cases [10, 13]. Dengue has become one of the most important health security risks in Pakistan, causing more outbreaks and having a higher mortality rate as compared to other vector-borne diseases.

Commercially, there is no antiviral drug or effective vaccine for dengue; the one licensed vaccine Dengvaxia (CYD-TDV) [14, 15] demonstrates an efficacy of 65.5% for ages 9 and above, and 44.6% for those below 9.[14]. Therefore, dengue control mainly relies on the control of mosquito vectors [16–18]. Traditional vector control strategies based on the use of insecticides are failing due to insecticide resistance in the vectors [19–21] and related environmental concerns [18, 19]. Consequently, alternative strategies have been investigated, including chemical or radiation-based sterile insect technique (SIT), gene alternation approaches such as RIDL (<u>R</u>elease of Insects carrying a <u>D</u>ominant <u>L</u>ethal Gene) and Incompatible Insect <u>T</u>echnique (IIT), a biological control method based on endosymbiont *Wolbachia* [16, 22, 23]. Incompatible insect technique, similar to SIT, focuses on release of *Wolbachia*-infected mosquitoes for population suppression [16, 24]. However, IIT holds an advantage due to specific *Wolbachia* strains that can block pathogen transmission, facilitate population modification, and manipulate the host’s reproductive mechanism [24]. <u>C</u>ytoplasmic Incompatibility (CI) is the most common reproductive manipulation induced by *Wolbachia,* causes sperm-egg incompatibility, leading to early embryonic death [22, 25]. *Wolbachia* is a maternally-transmitted obligate intracellular parasite found in at least 60% of insects [22]. Many VBDs like dengue, Zika, yellow fever, West Nile, chikungunya and malaria can be managed through *Wolbachia*-based control methods [20, 26].

*Ae. albopictus* naturally carries two strains of *Wolbachia* (*w*AlbA and *w*AlbB) [16], this phenomenon is known as superinfection and is common in insects [27, 28]. There are only a few reports of *Wolbachia* occurrence in *Ae. aegypti*, where *Wolbachia* presence was detected using modified and advanced techniques, revealing a similar infection as that of *w*AlbB in *Ae. albopictus* [29–31]. Naturally occurring *Wolbachia* strains in disease carrying mosquitoes exhibit limited resistance to virus transmission compared to artificially introduced strains [32, 33].

*Wolbachia* strains *w*Mel and *w*MelPop from *Drosophila* [5, 20] *w*Pip from *Culex pipiens* [34, 35] and *w*AlbB from *Ae. albopictus* are routinely used for artificial infection of *Ae. aegypti* [36, 37], and *Ae. albopictus* [38]. In *Ae. albopictus*, IIT is recommended, involving the release of incompatible males to reduce mean female fertility in the population [34, 35]. Population replacement requires introduction of an additional incompatible *Wolbachia* strain alongside the two natural strains already present in *Ae. albopictus* [39]. However, a triple infected *Ae. albopictus* population has proven unstable. For *Ae. aegypti*, predominantly uninfected in nature, both IIT and population replacement strategies are effective. IIT can be employed by creating a population artificially infected with *w*AlbB or *w*Mel strains and periodically releasing only-male infected mosquitoes [40]. Population replacement requires release of both male and female *Ae. aegypti*, and this self-sustaining strategy can operate without further releases if adequately monitored [41].

To investigate the presence of natural Wolbachia infection in *Ae. aegypti* and *Ae. albopictus*, we used marker gene-based metagenomic sequencing, a sensitive and reliable tool for detection of unculturable obligatory intracellular parasites like *Wolbachia* [42, 43]. We chose the most commonly used short-read MiSeq platform of Illumina which prefer to use V3/V4 hypervariable regions of 16s RNA gene [44].

## Methods

### Collection of Aedes mosquitoes

Adult mosquitoes were collected between March 2020 to November 2020 using Gravitraps, in seven different geographical regions in Peshawar district (Figure. 1). Further information about collection sites and collected samples has been listed in metadata (Table 1) and (Table S1). Adult *Aedes* were identified at the species level in the microscopic unit of the Department of Entomology, University of Agriculture Peshawar, using the morphological key of Rueda [45].

**Figure 1.**
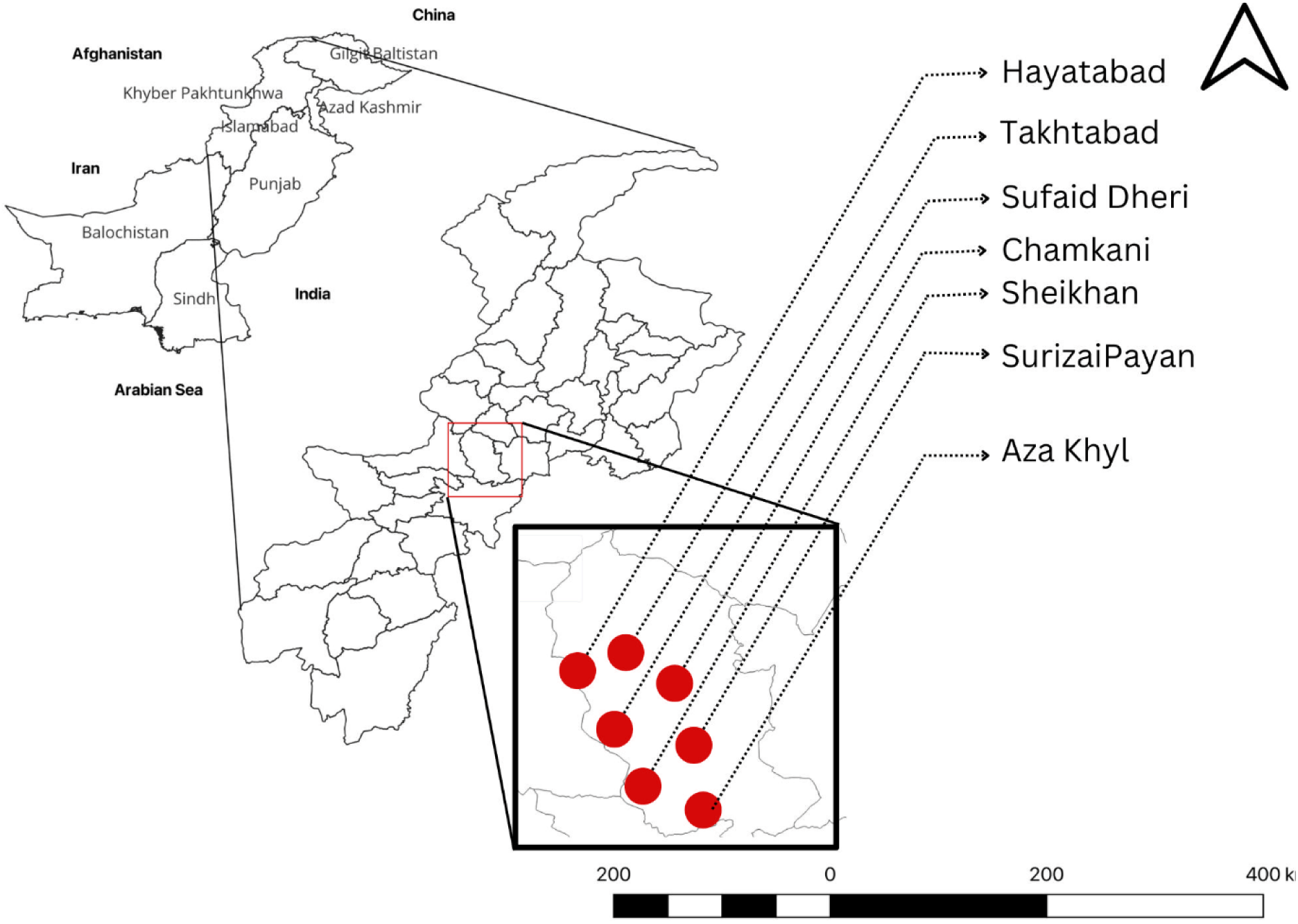
Collection sites. Samples were collected from six different neighborhoods within the Peshawar division. The first map displays the geographic location of Pakistan, while the second map focuses on the zoomed-in view of the Khyber Pakhtoonkhwa province. The Peshawar division is enclosed within a square, and collection sites are indicated by red dots on the zoomed-in map. The scale refers to the map of Khyber Pakhtoonkhwa. Source: source software QGIS version 3.28.3 (GNU General Public License), developed by the Open-Source Geospatial Foundation Project (http://qgis.org).

**Table 1.** Ae. aegypti and Ae. albopictus metadata.

|  | *Ae. Aegypti* | *Ae. albopictus* |
| --- | --- | --- |
| Taxon ID | 7159 | 7160 |
| Collection Time | 03/2020-09/2020 | 03/2020-09/2020 |
| City Latitude | 34.0149748 | 34.0149748 |
| City Longitude | 71.5804899 | 71.5804899 |
| Env Biome | ENVO:00000073,<br>ENVO:00000011 | ENVO:00000073,<br>ENVO:00000011 |
| Env Feature | ENVO:00000856 | ENVO:00000856 |
| Env Material | ENVO_00002005 | ENVO_00002005 |
| Sample Life Stage | Adult | Adult |
| Sample Sex | Female | Female |
| Sequencing Platform | Illumina | Illumina |

Male and female mosquitoes were identified by examining mouthparts and antennae. We separated 20 females each of *Ae. aegypti* and *Ae. albopictus* for DNA extraction.

### DNA extraction from female Ae. aegypti and Ae. albopictus

Before DNA extraction, samples were frozen in 100% ethanol at −20°C. Each mosquito was surface sterilized for 15 minutes with 70% ethanol before being washed in sterile distilled water.

Individual mosquitoes were placed in 1.5 mL Eppendorf tubes and were crushed and homogenized into a uniform powder. The DNA of each mosquito was extracted individually following manufacturer protocol (EZbioscience® DNA kit, https://www.ezbioscience.com/index.php/). Two separate DNA pools were created for metagenomic studies: each of 10 *Ae. aegypti* and 10 *Ae. albopictus* DNA. Each pool contained 2 µl of DNA extracted from each of 10 individual mosquitoes. In addition, the DNA of *Ae. albopictus* and *Ae. aegypti* was processed individually for DNA barcoding using COI (mitochondrial Cytochrome C oxidase I) gene. DNA quality testing was done using Nanodrop (Thermo Scientific, USA). Extracted DNA was stored at − 80°C for further use.

### Molecular characterization of COI gene by Sanger sequencing

After morphological identification, the identity was further confirmed through molecular characterization using species-specific primers followed by Sanger sequencing. A 710-bp fragment of the mitochondrial gene; cytochrome oxidase subunit 1 was amplified using a set of forward LCO-1490 (5’ GTCAACAAATCATAAAGATATTGG 3’) and reverse HCO-2198 (5’ TAA ACT TCA GGG TGA CCA AAA AAT CA 3’) primers [46]. The PCR reaction contained 2x KAPA HiFi HotStart ReadyMix and 1 µM of each primer (Forward/reverse). Amplification was performed in a Veriti thermal cycler with an initial denaturation at 95°C for 5 minutes, then 35 cycles at 95°C for 45 seconds (cyclic denaturation); 54°C for 45 seconds (annealing), and 72°C for 50 seconds (cyclic extension); followed by final elongation step at 72°C for 7 minutes. The PCR reaction was subjected to 1.5% agarose gel for 30 minutes to resolve the amplified product and visualize bands. The PCR products were sent to Macrogen services in South Korea for Sanger sequencing, utilizing the Applied Biosystems platform. The quality control of raw sequence data was done using FastQC to identify low-quality reads. Trimmomatic software was employed to remove bases with Phred scores below 20. The resulting high-quality sequences were generated in FASTA format for further analysis.

### Analysis of COI gene sequences

The obtained sequences were subjected to NCBI blast tools for the identification of homology patterns. Top identified and randomly selected sequences were aligned using the MUSCLE algorithm, in MEGA-X (Molecular Evolutionary Genetics Analysis) to determine phylogenetic relationships by neighbor-joining method. Bootstrap percentage values were obtained from 1000 replicates. Sequences with high similarity scores were downloaded from the NCBI database (http://blast.ncbi.nlm. nih.gov/Blast.cgi).

### Metagenomic sequencing of 16S rRNA using MiSeq

Pool DNA of *Ae. aegypti* and *Ae. albopictus* was used for metagenomic sequencing. Paired end reads from the V3/V4 region of 16S rRNA were generated on the Illumina MiSeq platform following the standard protocol (Macrogen; South Korea). The primers used in the study were 518F (5’ CCAGCAGCCGCGGTAATACG 3’) and 800R (5’ TACCAGGGTATCTAATCC 3’) [47]. Both forward and reverse primers were tagged with Illumina adapter, pad and linker sequences. The validated PCR libraries were used for sequencing on the Illumina MiSeq Platform. Raw FASTQ sequence reads were stored on the Illumina BaseSpace server for further analysis.

### Analysis of 16S metadata

The quality control check was performed on raw FASTQ files obtained from the Illumina BaseSpace website and analyzed with the Quantitative Insights into Microbial Ecology (QIIME) pipeline [48]. Sequences were trimmed to obtain a quality score less than PHRED 20. Low-quality, non-target, and chimeric reads were discarded and only valid reads were used for subsequent microbiome analyses. The reads with lengths between 350 bp to 550 bp were clustered into operational taxonomic units (OTU) using open reference OTU-picking against EZbiocloud 16S database version PKSSU4.0 [49]. Each read is identified at the species level with a 97% similarity cutoff. The reads below this cutoff were compiled and UCLUST [50] was used for *de novo* clustering to generate additional OTUs. The taxonomic hierarchy from phylum to species level was determined to measure the diversity of bacteria in *Aedes* microbiota. Alpha diversity indices ACE, Chao1, Jackknife, Simpson evenness and Shannon values were estimated based on the number and pattern of OTUs. Good’s coverage of the library was calculated based on the number of OTUs observed only once compared to the total number of OTUs present.

### Phylogenetic analysis of Wolbachia

Identified sequences of *Wolbachia* were subjected to BLAST, the resulting subject sequences were aligned using the MUSCLE (MUltiple Sequence Comparison by Log Expectation) algorithm, in MEGA X software (Molecular Evolutionary Genetics Analysis). The tree was constructed through the Neighbor-Joining (NJ) method to determine phylogenetic relationships between the sequences. Nodes represent hypothetical ancestors and arms represent species.

Sequences were downloaded from the NCBI database (http://blast.ncbi.nlm.nih.gov/Blast.cgi).

## Results

### Species Identification and Molecular Diversity of Ae. aegypti and Ae. albopictus

A total of 223 *Aedes* mosquitoes were collected. After morphological identification, 41 specimens were identified as female *Ae. aegypti*, while 68 were determined to be female *Ae. albopictus* (Figure. 2). The molecular identification through COI analysis performed on ten individual *Ae. albopictus* and *Ae. aegypti* mosquitoes resulted in successful identification of four *Ae. albopictus* and three *Ae. aegypti* samples. The remaining samples were either lost during the handling, transfer or their DNA quality was too low to be exactly identified. The neighbor-joining of COI sequences shows *Ae. aegypti* KPK4 (ON954492) and *Ae. aegypti* KPK2 (ON954493) grouped on external nodes with *Ae. aegypti* COI sequences associated with Faisalabad (KF406395) Pakistan, an indication of a distant common ancestor of the associated species (Figure. 3). Three haplotypes (HapI, HapII and HapIV) were found in association with KPK2 and KPK4 sequences. All three haplotypes have been reported in New Caledonia; this and our current report combined can predict their frequent occurrence in the tropics. *Ae. aegypti* KPK2 (ON954585) occupied a separate branch with an internal node occupied by COI sequences reported from Lahore (KF406395) and the closest external node occupied by sequences reported from Peshawar (KF406388), an indication of the closest common ancestor and high similarity in genetic makeup. *Ae. albopictus* sequences KPK1 (OQ998911) and KPK5 (OQ996886) were grouped on internal nodes with *Ae. albopictus* KPK5 (OQ996887) and *Ae. albopictus* KPK7 (OQ996888) occupying external nodes on the same monophyletic branch (Figure. 4). The group shows a distant phylogenetic relationship with CO1 sequences reported previously by NIBGE from Pakistan (Table S2), and sequences reported from China. The complete dataset can be explored using the provided GenBank accession numbers: ON954492 KPK4, ON954585 KPK3, ON954493 KPK2, OQ996886 KPK5, OQ996887 KPK6, OQ996888 KPK7, OQ998911 KPK1 16S rRNA metadata analysis MiSeq sequencing of the 16S rRNA amplicons generated 71,859 sequences in *Ae. aegypti* and 75,117 in *Ae. albopictus*. After the removal of the low-quality, non-target and chimeric reads, the number of valid reads was reduced to 65,457 in *Ae. aegypti* and 70,863 in *Ae. albopictus*. Little difference was observed between the average read lengths of *Ae. aegypti* (419 bp) and *Ae. albopictus* (418bp). A total of 62,749 (95.9%) reads in *Ae. aegypti* and 66,501 (93.8%) in *Ae. albopictus* were successfully identified at the species level which leads to the successful identification of 239 and 921 species in *Ae. aegypti* and *Ae. albopictus* respectively using reference database. With a 97% similarity cutoff, the number of OTUs observed in *Ae. aegypti* (259) and *Ae. albopictus* (1002) was greater than the actual number of identified species in the respective microbiota of the two species, an indication that OTU count doesn’t necessarily equate to the actual number of identified species (Table 2).

**Figure 2:**
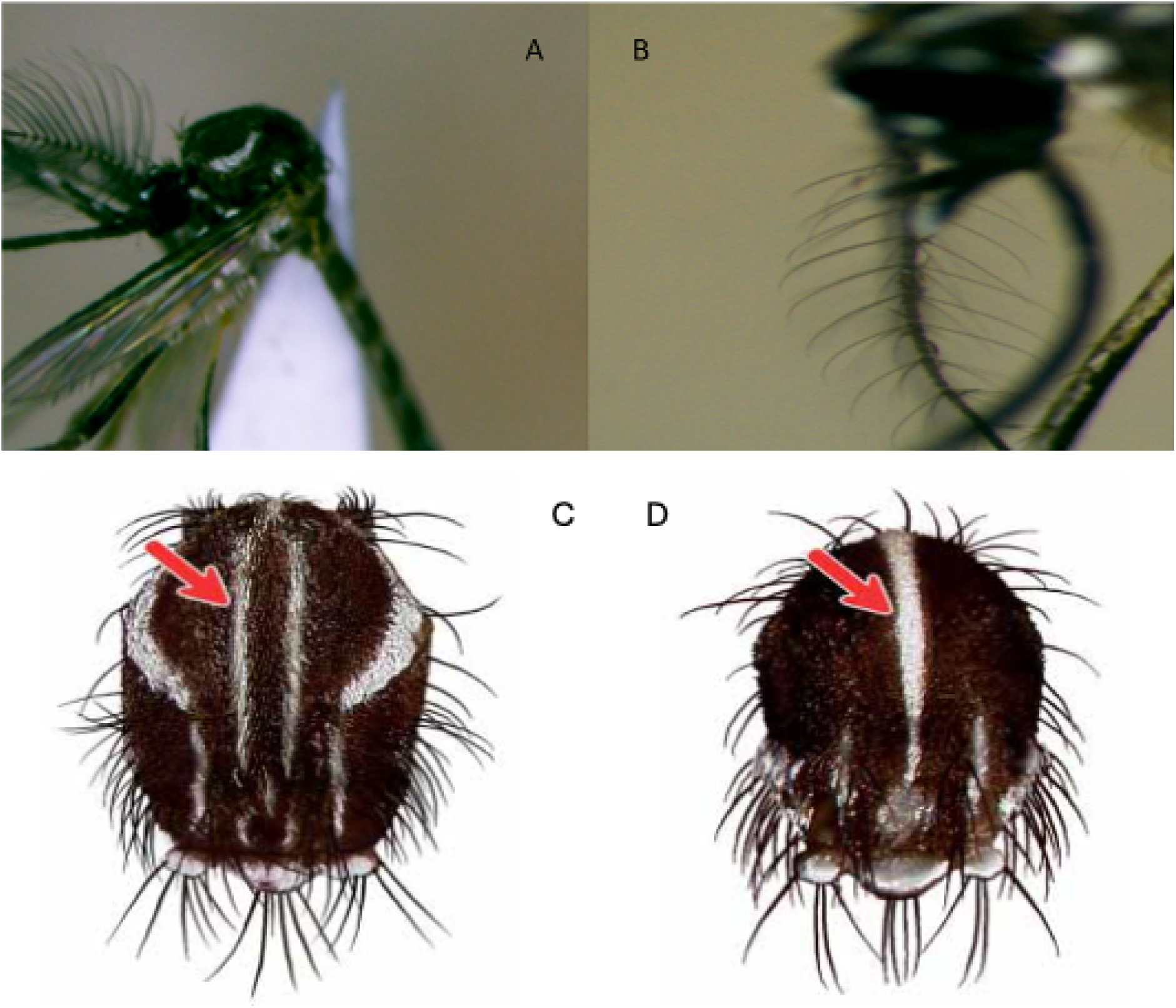
Morphological identification. A. Head and thorax of a male *Ae. aegypti* mosquito. B: Head and thorax of a female *Ae. aegypti mosquito*. Differences in the antennae can be seen. C. Identification marks on *Ae. aegypti* head region D. Identification of *Ae. albopictus.* (Morphological key and Figure 2 C and D are modified from Rueda 2004)

**Figure 3.**
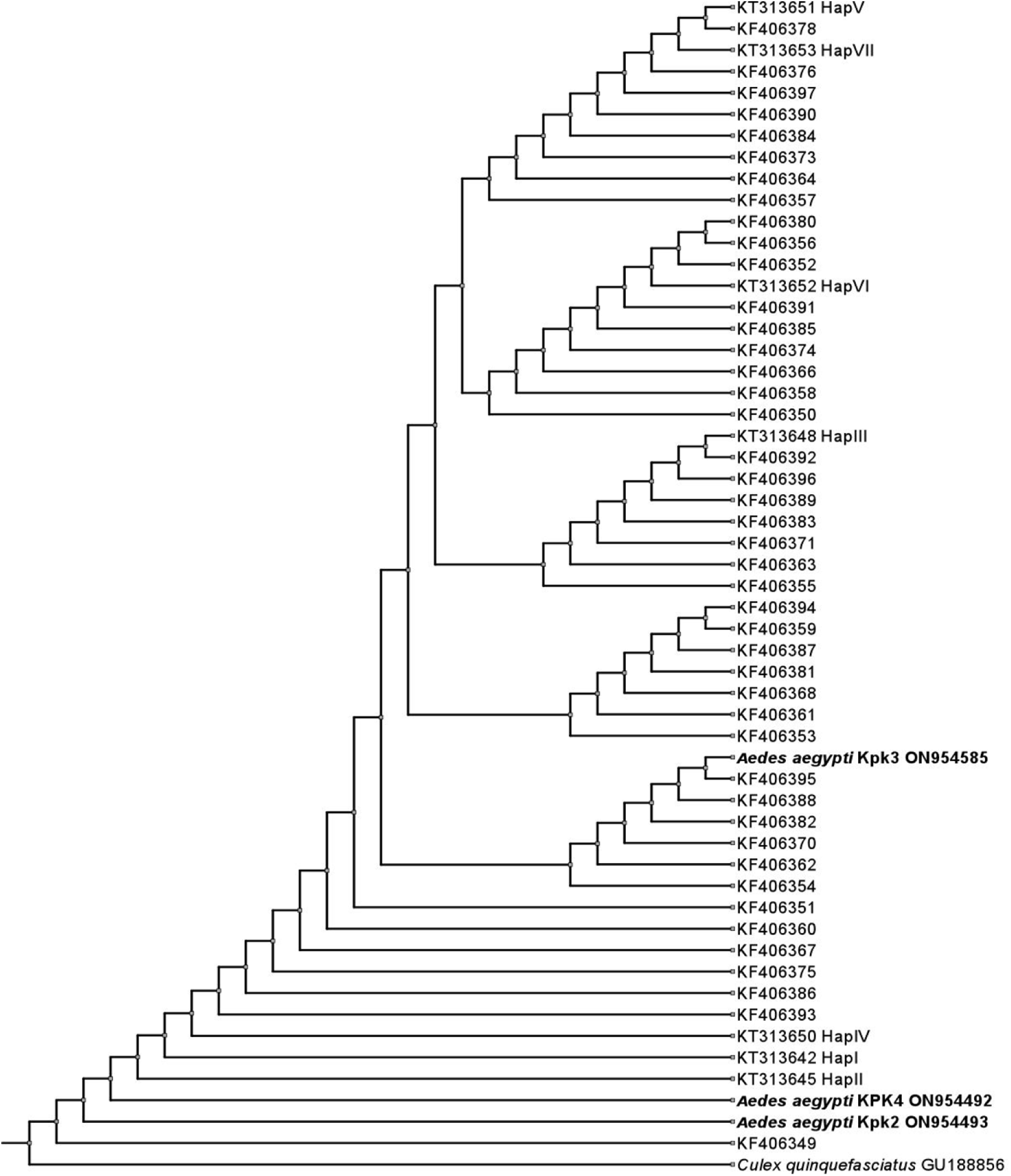
*Aedes aegypti* neighbor-joining (NJ) distance tree based on COI gene sequences. Phylogenetic tree was constructed using MEGA X software. Sequences are aligned using MUSCLE algorithm. The tree shows evolutionary relationship of mitochondrial COI genes of identified *Aedes aegypti* with COI genes reported previously from other studies. COI genes in tree are represented by GenBank accession numbers. *Culex quinquefasciatus* is taken as outgroup in the phylogenetic tree.

**Figure 4:**
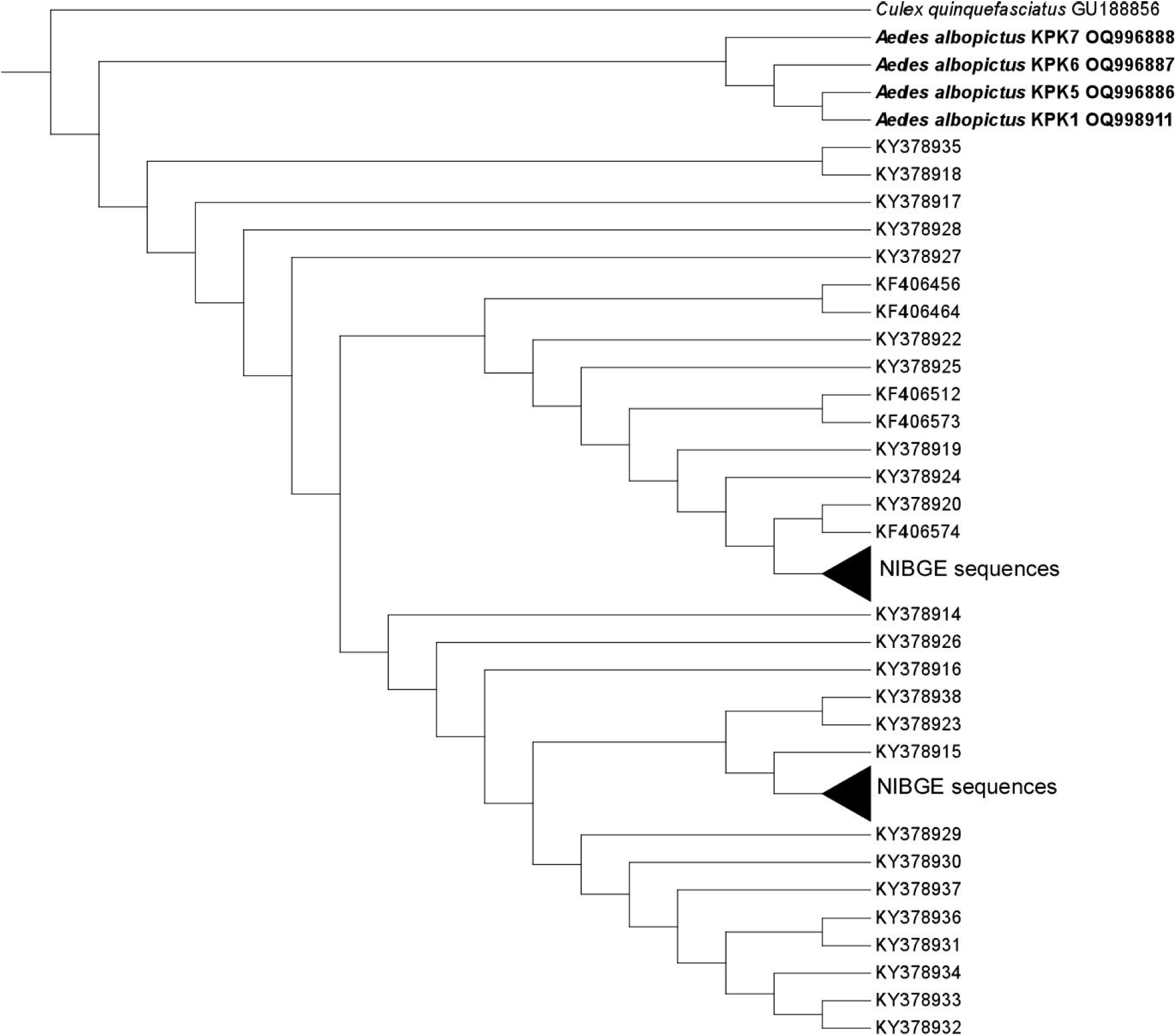
*Ae. albopictus* (NJ) distance tree based on COI gene sequences. Phylogenetic tree was constructed using MEGA X software. Sequences are aligned using MUSCLE algorithm. Sequences are represented by GenBank accession numbers. The accession numbers of CO1 sequences reported by NIBGE (National Institute of Biotechnology and Genetic Engineering) are listed in Supplementary Information Table S2. *Culex quinquefasciatus* is taken as outgroup in the tree.

**Table 2.** Sequenced reads, OTU count and number of microbial species detected in *Ae. aegypti* and *Ae. albopictus* microbiota.

|  | Total<br>Sequences | Valid Reads | Average Read<br>Length | OTU Count | Identified<br>Species |
| --- | --- | --- | --- | --- | --- |
| *Ae. aegypti* | 71,859 | 65,457 | 419bp | 259 | 239 |
| *Ae. albopictus* | 75,117 | 70,863 | 418bp | 1002 | 921 |

### Alpha diversity

Alpha diversity indices were measured using the number and pattern of OTUs observed in the sample. The species richness indicators Ace, Chao and Jackknife show higher values for *Ae. albopictus* as compared to *Ae. aegypti*, an indication of comparatively rich microbiota in *Ae. albopictus*. Species evenness through Shannon’s values were also observed higher for *Ae. albopictus*. The lower Simpson value in *Ae. albopictus* (0.03) as compared to *Ae. aegypti* (0.07) show comparatively greater diversity and an even distribution of species in *Ae. albopictus*.

Phylogenetic diversity was used as an additional species richness indicator; the values were again found higher in *Ae. albopictus* (1727) than *Ae. aegypti* (610). The Good’s coverage of library value was 100% for *Ae. aegypti* indicating additional rounds of sequencing may not find any new species. The 99.9% value in *Ae. albopictus* shows few unrecognized sequences in the sample (Table 3).

**Table 3.** Comparative analysis of alpha diversity between *Ae. aegypti* and *Ae. albopictus*.

|  | ACE | CHAO | Jackknife | Shannon | Simpson | Phylogenetic<br>Diversity |
| --- | --- | --- | --- | --- | --- | --- |
| *Ae. aegypti* | 269.71 | 269.2 | 277 | 3.35 | 0.07 | 610 |
| *Ae. Albopictus* | 1032.49 | 1057.88 | 1081.88 | 4.69 | 0.03 | 1727 |

**Table 4.**
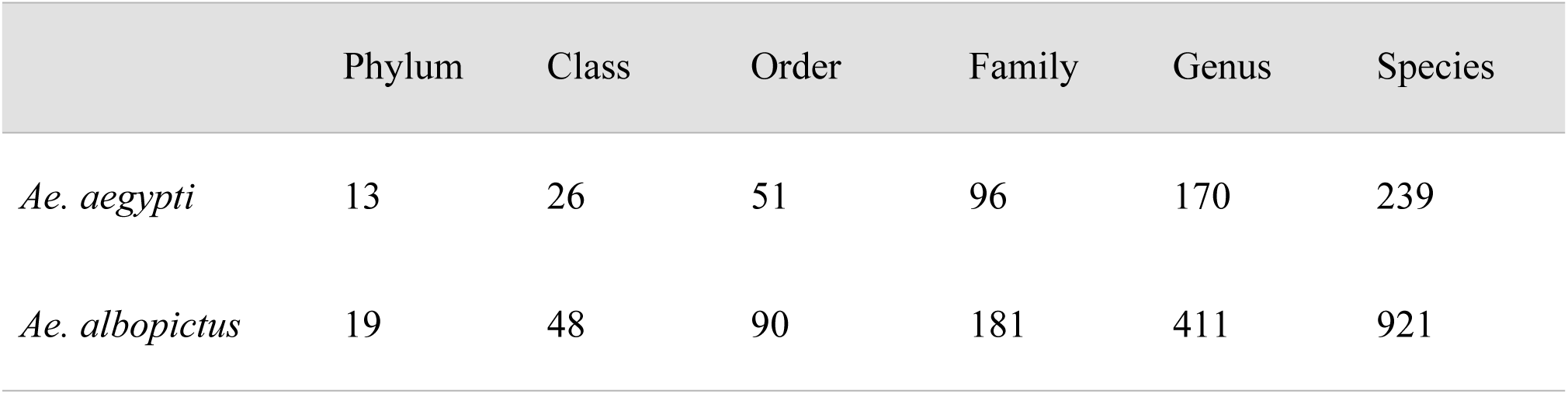
Comparative analysis of number of taxa detected at six different levels in *Ae. aegypti* and *Ae. albopictus* microbiota.

|  | Phylum | Class | Order | Family | Genus | Species |
| --- | --- | --- | --- | --- | --- | --- |
| *Ae. aegypti* | 13 | 26 | 51 | 96 | 170 | 239 |
| *Ae. albopictus* | 19 | 48 | 90 | 181 | 411 | 921 |

### Comparative analysis of microbial diversity

A total of 239 bacterial species were detected in *Ae. aegypti*, while 921 were detected in *Ae. albopictus*. Both species shared eleven bacterial groups at the phylum level (Figure S1). Phylum Proteobacteria was the most abundant group in both *Aedes* species with a relative abundance of 80.04% in *Ae. aegypti* and 52.33% in *Ae. albopictus* (Figure.5). Among other prominent phyla, Actinobacteria accounted for 10.39% and 4.99%, Firmicutes comprised 3.91% and 22.10%, and Bacteroidetes represented 18.60% and 4.95% in *Ae. aegypti* and *Ae. albopictus*, respectively. The class Gammaproteobacteria has the highest concentration in both species with a 46.07% proportion in *Ae. aegypti* and 36.07% in *Ae. albopictus.* The other prominent shared classes between the two species were Betaproteobacteria, Alphaproteobacteria, Actinobacteria and Bacilli with relative abundances of (17.76%), (16.17%), (10.32%), (3.84%) in *Ae. aegypti*, and (4.55%), (11.53%), (21.83%), (15.43%) in *Ae. albopictus*, respectively. Order Enterobacterales was the most abundant group in both species with a 23.44% proportion in *Ae. aegypti* and 25.03% in *Ae. albopictus.* The other prominent detected orders in *Ae. aegypti* were Burkholderiales (16.12%), Pseudomonadales (11.10%), Ricketsiales (8.92%) and Corynebacteriales (8.84%), while in *Ae. albopictus,* the notable orders were Corynebacteriales (8.80%), Bacillales (7.58%), Lactobacillales (7.58%) and Pseudomonadales (7.45%). In *Ae. aegypti*, the genus *Wolbachia* ranked as the fourth most abundant genus, with *Wolbachia pipientis* representing the entirety of 8.92% of the total abundance. However, in *Ae. albopictus*, *Wolbachia* exhibited a relatively lower concentration of 3.71%, placing it as the ninth most abundant genus in the microbiota. We detected a co-infection of *Wolbachia bourtzisii* (2.37%) and *Wolbachia pipientis* (1.33%) within *Ae. albopictus* microbiota. Comprehensive data on the bacteriome, spanning from phylum to species level, can be found in the supplementary data (Table S3).

**Figure 5.**
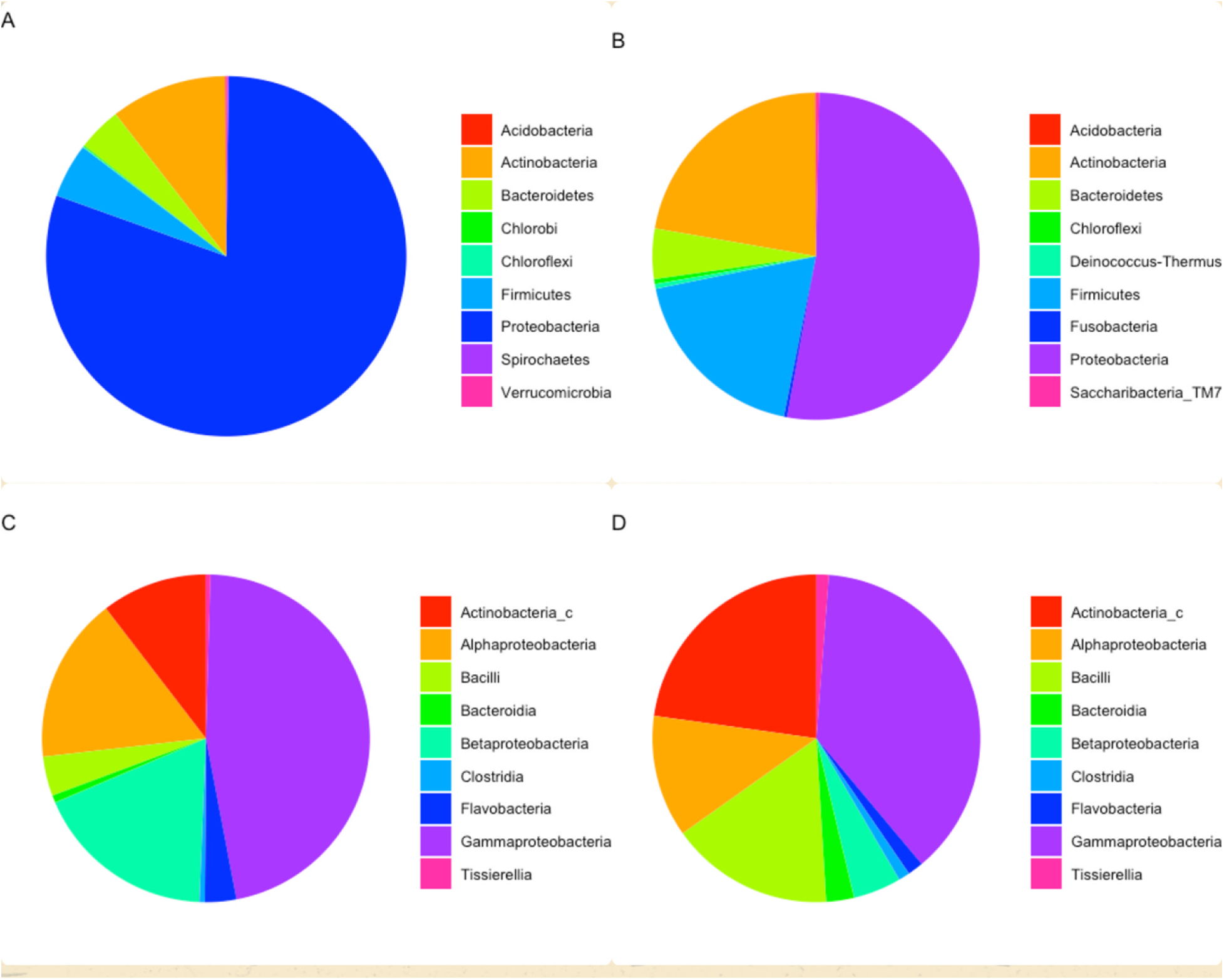
(A, B, C, D): Comparative analysis of microbial diversity between *Aedes aegypti* and *Aedes albopictus.* ‘A’ represents relative abundance of phyla in *Aedes aegypti*; ‘B’ represents relative abundance of phyla in *Aedes albopictus*; ‘C’ represents relative abundance of classes in *Aedes aegypti*; ‘D’ represents relative abundance of classes in *Aedes albopictus*; ‘

**Figure 5.**
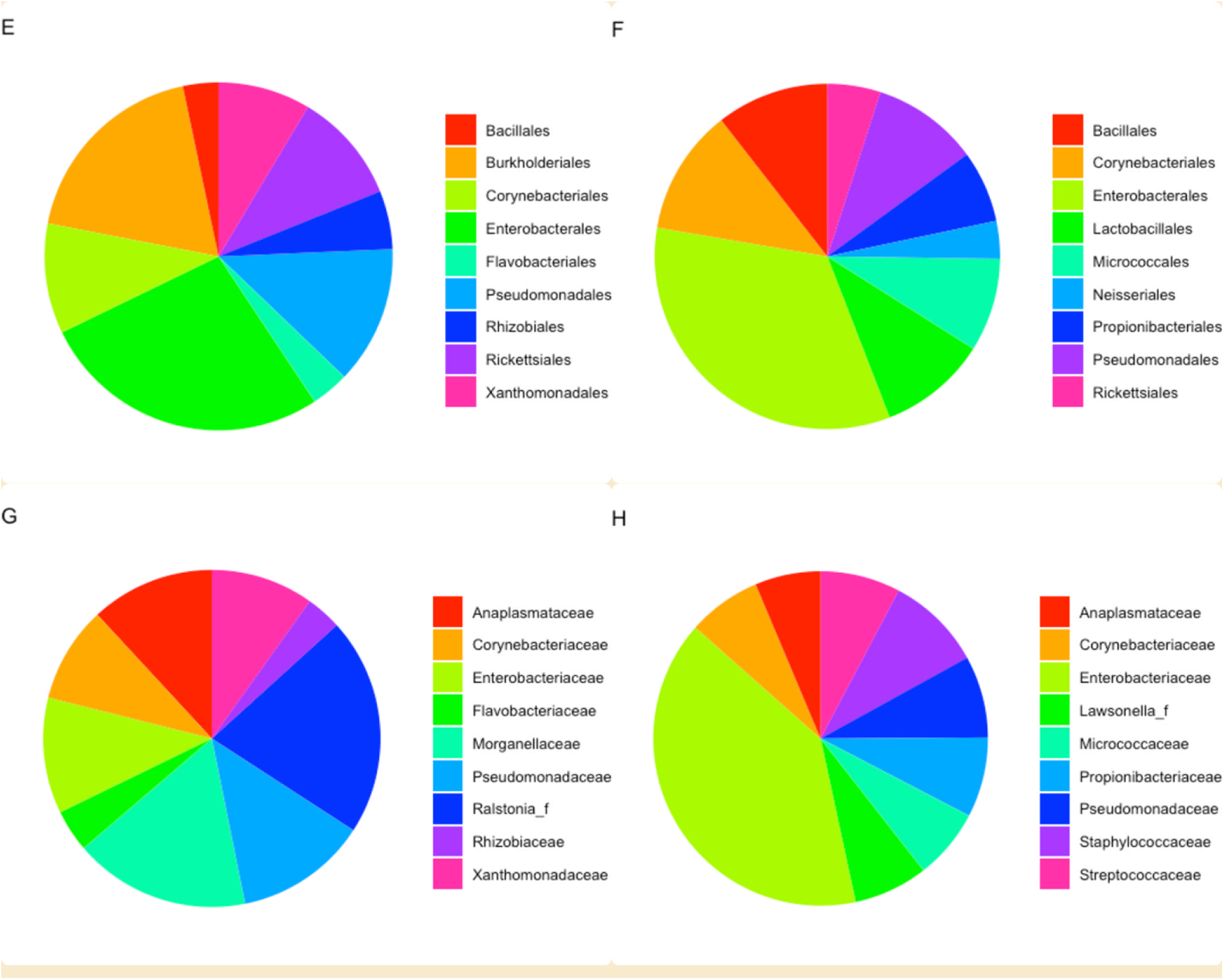
(E, F, G, H): Comparative analysis of microbial diversity between *Aedes aegypti* and *Aedes albopictus.* E’ represents relative abundance of orders in *Aedes aegypti*; ‘F’ represents relative abundance of orders in *Aedes albopictus*. ‘G’ represents relative abundance of families in *Aedes aegypti*; ‘H’ represents relative abundance of families in *Aedes albopictus*; ‘

**Figure 5.3.**
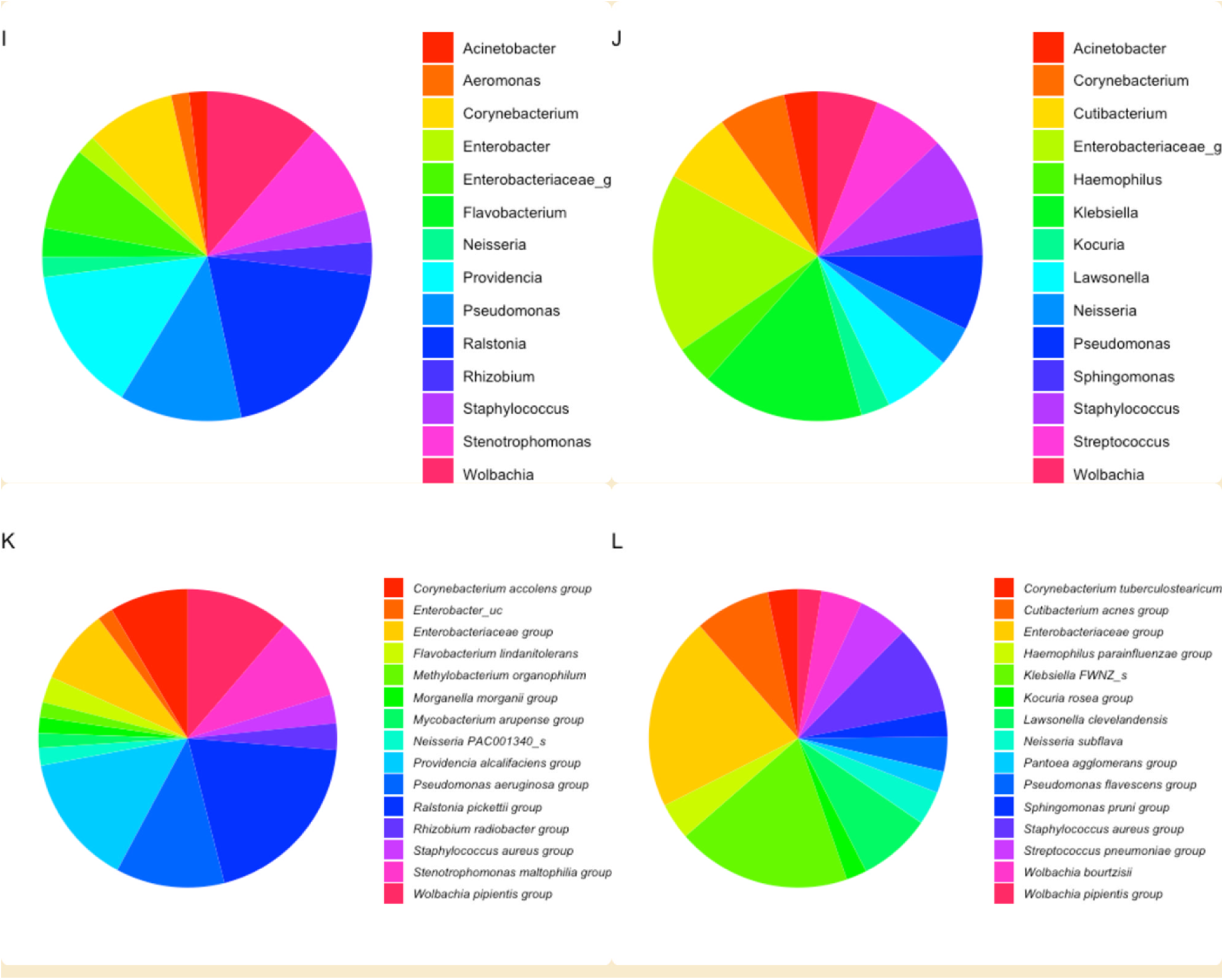
(I, J, K, L): Comparative analysis of microbial diversity between *Aedes aegypti* and *Aedes albopictus.* I’ represents relative abundance of genera in *Aedes aegypti*; ‘J’ represents relative abundance of genera in *Aedes albopictus*; ‘K’ represents relative abundance of species in *Aedes aegypti*; ‘L’ represents relative abundance of species in *Aedes albopictus*.

### Phylogenetic analysis of identified Wolbachia species

We tried to uncover the evolutionary relationship and genetic variations between identified *Wolbachia* strains in this study and previously reported *Wolbachia* strains by other investigators through phylogenetic analysis. The identified *Ae. aegypti* strains *w*PipAgy_PK2, *w*PipAgy_PK3, and *Ae. albopictus* strain *w*PipAlb_PK4 are grouped on internal sister nodes with group B strains *w*Agi (MF999263) *Ae. aegypti*, *w*No (CP003883.1) *Drosophila simulans*, *w*Pip (AM999887.1) *Cx. quinquefasciatus*, *w*AlbB (KX155506.1) *Ae. albopictus*, *w*CPip (X61768) *Culex pipiens*, *w*Nasovit (M84686) Wasp *Nasonia*. This shows a divergence from the closest same ancestor, indicating high similarities in the strains obtained from these otherwise different insect species. *Wolbachia* strains *w*PipAlb_PK1, *w*PipAlb_PK5, *w*PipAlb_PK6, and *w*PipAlb_PK7 belonged to *Ae. albopictus*. This cluster does not have sister groups, indicating unique genetic makeup and distant phylogenetic relationship with other groups (Figure. 6).

**Figure 6:**
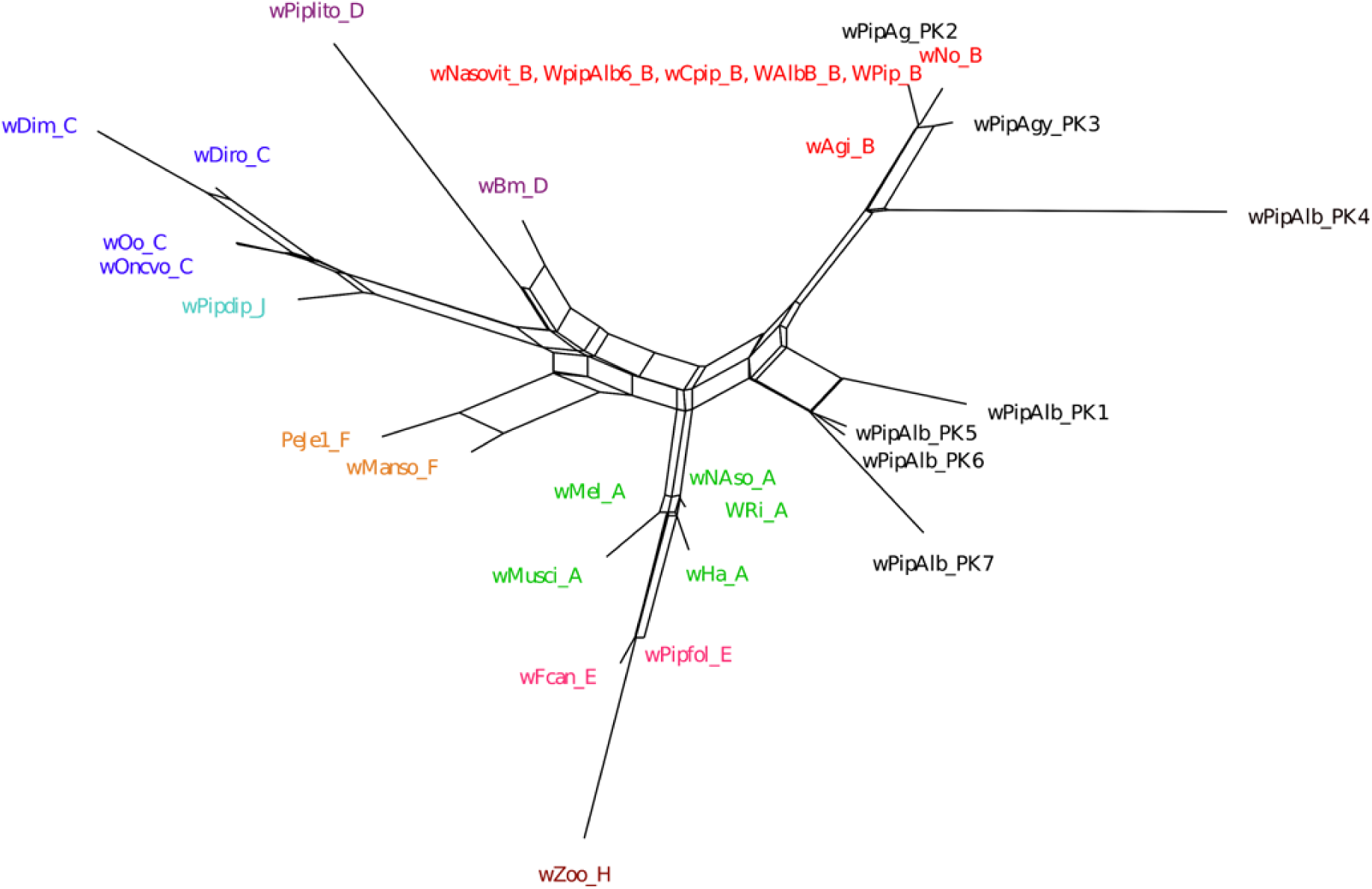
Phylogenetic analysis of *Wolbachia*. Tree was constructed by neighbor joining method based on 16S rRNA sequences in MEGA X software. Sequences are aligned using the MUSCLE program. Bootstrap percentage values were obtained from 1000 replicates. GenBank accession numbers of sequences used in analyses are shown on the phylogenetic tree. Closely related *Wolbachia* Sequences are shown in eight supergroups, represented by different colors in the tree.

## Discussion

The density of *Wolbachia* within insect hosts is subject to the intricate interplay of multiple factors, including the host’s species, the specific strain of the *Wolbachia* bacterium, as well as environmental variables such as habitat, temperature, and early growth-stage food availability [51]. We employed a metagenomics approach to identify intracellular bacteria within the vectors responsible for transmitting dengue, Zika, chikungunya, and yellow fever viruses. Our findings revealed the presence of *Wolbachia* in both mosquito vectors, with an intriguing discovery of greater bacterial diversity in *Ae. albopictus* compared to *Ae. aegypti*.

*Ae. albopictus* is abundant in Peshawar [52, 53], whereas *Ae. aegypti* is sporadically present [52]. We collected *Ae. albopictus* from five out of seven collection sites while *Ae. aegypti* was only collected from three areas: Sheikhan, Hayatabad and Sufaid Dheri, located near Khyber district in the west of Peshawar, where *Ae. aegypti* is reportedly endemic [53]. This raises concerns about the vector’s potential expansion beyond its boundaries to other regions. The COI sequences identified in *Ae. aegypti* and *Ae. albopictus* displayed a close resemblance with the *Ae.* sequences reported previously from various cities of Pakistan [54]. *Ae. aegypti* KPK4, and *Ae. aegypti* KPK2 showed close phylogenetic relationship with haplotypes from the Pacific [55]. This finding suggests a potential genetic connection between *Ae. aegypti* populations in Pakistan and the Pacific region. We detected a total of 239 bacterial species in *Ae. aegypti*, while in *Ae. albopictus* we were able to detect 921 species. At the phylum level, both species shared four groups, with phylum Proteobacteria prevailing as the most abundant group among them. These findings were in accordance with the previously reported data from *Ae. aegypti* and *Ae. albopictus* [56, 57]. The other major detected phyla were Actinobacteria, Firmicutes and Bacteroidetes. Phylum Bacteroidetes have been detected in *Ae. aegypti* larvae and water [30], indicating habitat of a mosquito has a great impact on its microbiota. The shared classes between the two *Ae.* species were gram-negative bacteria Gammaproteobacteria, Alphaproteobacteria, Flavobacteria, Betaproteobacteria and gram-positive bacteria Actinobacteria. These classes get frequently associated with *Aedes* mosquitoes, suggesting that these may be the most common bacterial groups associated with *Aedes* microbiota.

We found several bacteria that are commonly associated with insect vectors *Culex*, *Anopheles* and *Aedes* [56, 58–61], *Anopheles darlingi* [62] *An. albimanus* [63], *Culicoides sonorensis* [64], *Culex quinquefasciatus* [65]. Our study also identified several bacterial groups that are commonly associated with soil [66] suggesting habitat and breeding places interaction with soil, which is a rich source of microorganisms. The Gammaproteobacteria is an important core taxon in the guts of soil invertebrates and also acts as a potential indicator of soil pollutants in these invertebrates [67]. Family Enterobacteriaceae emerged as a prominent group in both *Ae. aegypti* and *Ae. albopictus* microbiota. Notably, several members of this family reported in our data are suggested to provide an additional nitrogen source in fruit fly *Ceratitis capitata* [68]. Several members of the genus *Bacillus* detected in both *Aedes* species are thought to be involved in cellulose and hemicellulose degradation in termites [69], and malathion and permethrin insecticides degradation [57]. The *Serratia marcescens* species from genus *Serratia* was detected in a small proportion (0.4%) within *Ae. aegypti* microbiota which is known to enhance arboviruses susceptibility in *Ae. aegypti* [70].

Some significant differences were noted between the bacterial composition of the two *Aedes* species. The *Ae. albopictus* microbiota showed comparatively higher diversity suggesting a more evolved commensalism with bacterial species. This was in contrast with the previous findings where bacterial diversity of *Ae. aegypti* was found comparatively higher than *Ae. albopictus* [58]. The differences in the microbial composition can be attributed to the factors like genetic differences, water and habitat conditions and the impact of seasonal and climate changes. Environmental changes throughout collection greatly impact the types of microbes acquired by the insects in the same habitat [30, 71].

In Pakistan, a study conducted in 2022 confirmed the presence of *Wolbachia* infection in *Ae. albopictus* [72]. However, despite efforts, *Wolbachia’s* presence could not be detected in the principal dengue vector *Ae. aegypti* [73]. Using a metagenomic approach, we were able to detect *Wolbachia* infection in both dengue virus vectors. In *Ae. aegypti,* we detected *Wolbachia pipientis* infection while co-infection of *Wolbachia pipientis* and *Wolbachia bourtzisii* was detected in *Ae. albopictus*. *Wolbachia pipientis* is a commonly found insect endosymbiont and has been explored extensively by investigators. Several investigators have detected *Wolbachia pipientis* in the microbiota of *Ae. albopictus* [74, 75] while few have detected in *Ae. aegypti* [27, 29]. The *Wolbachia bourtzisii* has been detected in *Ae. albopictus* [76], *Drosophila melanogaster* [77], *Hemiptera* families *Dactylopiidae* (*Dactylopius coccus*) [78] and *Lygaeidae* (*Oxycarenus laetus*) [79]. In the case of confirmed natural *Wolbachia pipientis* infection in *Ae. aegypti*, an effective IIT-based control program will require the release of males carrying *Wolbachia bourtzisii* strains, such as *w*Mel. Our results also provide insight into the bacterial composition of *Aedes* microbiota. The symbioses between insects and vertically transmitted endosymbiotic bacteria can have a major influence on insect physiology and behavior [80, 81] which in turn can impact *Wolbachia*-mediated DENV blocking in dengue vectors.

### Conclusions

The presence of both *Ae. aegypti* and *Ae. albopictus* pose a significant threat to public health in Peshawar, Khyber Pakhtoonkhwa, Pakistan. However, the low dengue virus transmission can be attributed to the presence of endosymbiont bacteria *Wolbachia* in both vector species. Among bacterial phyla found in *Aedes*, eight were unique to *Ae. albopictus*, two were unique to *Ae. aegypti* while eleven phyla were shared between the two species. These findings show that *Ae. albopictus* harbors greater bacterial diversity compared to *Ae. aegypti*.

## Ethical approval and consent to participate

Not applicable.

## Consent for publication

Not applicable.

## Acknowledgments

The research was financially supported by Mr. Mark Drew, Executive Director of the Australian-Khyber Pakhtunkhwa Agricultural Development Organization (AKPADO), Brisbane QLD, Australia.

## Appendix A. Supplementary material

Supplementary file 1 Table S1

Collection Sites coordinates, Time period, and Collected Specimens.

Supplementary file 2 Table S2

Accession numbers and depositor information of CO1 sequences used in Figure 3.

Supplementary file 3 Figure S1

The number of shared and unique taxa in *Ae. aegypti* and *Ae. albopictus* at the phylum level

Supplementary file 4 Table S3

Complete microbiota of *Ae. aegypti* and *Ae. albopictus*.

## Authorship contribution statement

GN and MF conceived and designed the study and GN performed all the experimental work. RuR and FH helped with data analysis and reviewing the manuscript. IK provided mosquitoes samples and helped designed the study. GN wrote the initial draft and finalized it with the supervision of MF and RuR. All authors read and approved the final manuscript.

## Data availability

Data supporting the conclusions of this study are available in the GenBank repository (https://www.ncbi.nlm.nih.gov/genbank/). The given accession numbers to the COI sequences are; ON954492 KPK4, ON954585 KPK3, ON954493 KPK2, OQ996886 KPK5, OQ996887 KPK6, OQ996888 KPK7, OQ998911 KPK1. Metagenome data is available in BioSample Submission Portal as Bio-Project PRJNA933924.

## References

1. Wilson AL, Courtenay O, Kelly-Hope LA, Scott TW, Takken W, Torr SJ, et al. The importance of vector control for the control and elimination of vector-borne diseases. PLoS Negl Trop Dis. 2020;14(1):e0007831. doi: 10.1371/journal.pntd.0007831. PubMed PMID: 31945061; PubMed Central PMCID: PMCPMC6964823.

2. WH Organization, UNICEF. Global vector control response 2017-2030. 2017.

3. Coates SJ, Norton SA. The effects of climate change on infectious diseases with cutaneous manifestations. International journal of women’s dermatology. 2021;7(1):8–16.

4. Choumet V, Despres P. Dengue and other flavivirus infections. Rev Sci Tech. 2015;34(2):473-8, 67–72. PubMed PMID: 26601449.

5. Dorigatti I, McCormack C, Nedjati-Gilani G, Ferguson NM. Using Wolbachia for dengue control: insights from modelling. Trends in parasitology. 2018;34(2):102–13.

6. Brito AF, Machado LC, Oidtman RJ, Siconelli MJL, Tran QM, Fauver JR, et al. Lying in wait: the resurgence of dengue virus after the Zika epidemic in Brazil. Nature Communications. 2021;12(1):1–13.

7. Chan Y, Tan H, Seah C, Li J, Chow V, Salahuddin N, et al. Dengue haemorrhagic fever outbreak in Karachi, Pakistan, 1994. 1995.

8. Khan J, Khan I, Amin I. A comprehensive entomological, serological and molecular study of 2013 dengue outbreak of Swat, Khyber Pakhtunkhwa, Pakistan. PLoS One. 2016;11(2):e0147416.

9. Khan E, Kisat M, Khan N, Nasir A, Ayub S, Hasan R. Demographic and clinical features of dengue fever in Pakistan from 2003–2007: a retrospective cross-sectional study. PloS one. 2010;5(9):e12505.

10. Khan J, Ghaffar A, Khan SA. The changing epidemiological pattern of Dengue in Swat, Khyber Pakhtunkhwa. PloS one. 2018;13(4):e0195706.

11. Wesolowski A, Qureshi T, Boni MF, Sundsøy PR, Johansson MA, Rasheed SB, et al. Impact of human mobility on the emergence of dengue epidemics in Pakistan. Proceedings of the National Academy of Sciences. 2015;112(38):11887–92.

12. Rasheed S, Butlin R, Boots M. A review of dengue as an emerging disease in Pakistan. Public health. 2013;127(1):11–7.

13. Abdullah SA, Salman M, Din M, Khan K, Ahmad M, Khan FH, et al. Dengue outbreaks in Khyber Pakhtunkhwa (KPK), Pakistan in 2017: an integrated disease surveillance and response system (IDSRS)-based report. Polish journal of microbiology. 2019;68(1):115.

14. Hadinegoro SR, Arredondo-García JL, Capeding MR, Deseda C, Chotpitayasunondh T, Dietze R, et al. Efficacy and long-term safety of a dengue vaccine in regions of endemic disease. New England Journal of Medicine. 2015;373(13):1195–206.

15. Deen J. The dengue vaccine dilemma: balancing the individual and population risks and benefits. PLoS medicine. 2016;13(11):e1002182.

16. Zhang D, Zheng X, Xi Z, Bourtzis K, Gilles JR. Combining the sterile insect technique with the incompatible insect technique: I-impact of Wolbachia infection on the fitness of triple-and double-infected strains of Aedes albopictus. PloS one. 2015;10(4):e0121126.

17. Xu T-L, Han Y, Liu W, Pang X-Y, Zheng B, Zhang Y, et al. Antivirus effectiveness of ivermectin on dengue virus type 2 in Aedes albopictus. PLoS neglected tropical diseases. 2018;12(11):e0006934.

18. Achee N, Gould F, Perkins Jr T. RCR, Morrison AC, Ritchie SA, et al. A Critical Assessment of vector control for dengue prevention PLoS Negl Trop Dis. 2015;9(5):e0003655.

19. Epelboin Y, Chaney SC, Guidez A, Habchi-Hanriot N, Talaga S, Wang L, et al. Successes and failures of sixty years of vector control in French Guiana: what is the next step? Memórias do Instituto Oswaldo Cruz. 2018;113.

20. Walker T, Johnson P, Moreira L, Iturbe-Ormaetxe I, Frentiu F, McMeniman C, et al. The wMel Wolbachia strain blocks dengue and invades caged Aedes aegypti populations. Nature. 2011;476(7361):450-3.

21. Rahman RU, Souza B, Uddin I, Carrara L, Brito LP, Costa MM, et al. Insecticide resistance and underlying targets-site and metabolic mechanisms in Aedes aegypti and Aedes albopictus from Lahore, Pakistan. Scientific Reports. 2021;11(1):4555.

22. Achee NL, Grieco JP, Vatandoost H, Seixas G, Pinto J, Ching-Ng L, et al. Alternative strategies for mosquito-borne arbovirus control. PLoS neglected tropical diseases. 2019;13(1):e0006822.

23. Qsim M, Ashfaq UA, Yousaf MZ, Masoud MS, Rasul I, Noor N, et al. Genetically modified Aedes aegypti to control dengue: a review. Critical Reviews™ in Eukaryotic Gene Expression. 2017;27(4).

24. Wang GH, Gamez S, Raban RR, Marshall JM, Alphey L, Li M, et al. Combating mosquito-borne diseases using genetic control technologies. Nat Commun. 2021;12(1):4388. doi: 10.1038/s41467-021-24654-z. PubMed PMID: 34282149; PubMed Central PMCID: PMCPMC8290041 this arrangement have been reviewed and approved by the University of California, San Diego in accordance with its conflict of interest policies. All other authors declare no competing interests.

25. McMeniman CJ, Lane RV, Cass BN, Fong AW, Sidhu M, Wang Y-F, et al. Stable introduction of a life-shortening Wolbachia infection into the mosquito Aedes aegypti. Science. 2009;323(5910):141-4.

26. Shaw WR, Marcenac P, Childs LM, Buckee CO, Baldini F, Sawadogo SP, et al. Wolbachia infections in natural Anopheles populations affect egg laying and negatively correlate with Plasmodium development. Nat Commun. 2016;7:11772. doi: 10.1038/ncomms11772. PubMed PMID: 27243367; PubMed Central PMCID: PMCPMC4895022.

27. Teo C, Lim P, Voon K, Mak J. Detection of dengue viruses and Wolbachia in Aedes aegypti and Aedes albopictus larvae from four urban localities in Kuala Lumpur, Malaysia. Trop Biomed. 2017;34:583–97.

28. Zhou W, Rousset F, O’Neill S. Phylogeny and PCR–based classification of Wolbachia strains using wsp gene sequences. Proceedings of the Royal Society of London Series B: Biological Sciences. 1998;265(1395):509-15.

29. Kulkarni A, Yu W, Jiang J, Sanchez C, Karna AK, Martinez KJ, et al. Wolbachia pipientis occurs in Aedes aegypti populations in New Mexico and Florida, USA. Ecology and evolution. 2019;9(10):6148–56.

30. Coon KL, Brown MR, Strand MR. Mosquitoes host communities of bacteria that are essential for development but vary greatly between local habitats. Molecular ecology. 2016;25(22):5806–26.

31. Ross PA, Callahan AG, Yang Q, Jasper M, Arif MA, Afizah AN, et al. An elusive endosymbiont: Does Wolbachia occur naturally in Aedes aegypti? Ecology and Evolution. 2020;10(3):1581–91.

32. Glaser RL, Meola MA. The native Wolbachia endosymbionts of Drosophila melanogaster and Culex quinquefasciatus increase host resistance to West Nile virus infection. PLoS One. 2010;5(8):e11977. doi: 10.1371/journal.pone.0011977. PubMed PMID: 20700535; PubMed Central PMCID: PMCPMC2916829.

33. Mousson L, Zouache K, Arias-Goeta C, Raquin V, Mavingui P, Failloux AB. The native Wolbachia symbionts limit transmission of dengue virus in Aedes albopictus. PLoS Negl Trop Dis. 2012;6(12):e1989. doi: 10.1371/journal.pntd.0001989. PubMed PMID: 23301109; PubMed Central PMCID: PMCPMC3531523.

34. Calvitti M, Moretti R, Skidmore AR, Dobson SL. Wolbachia strain wPip yields a pattern of cytoplasmic incompatibility enhancing a Wolbachia-based suppression strategy against the disease vector Aedes albopictus. Parasit Vectors. 2012;5:254. doi: 10.1186/1756-3305-5-254. PubMed PMID: 23146564; PubMed Central PMCID: PMCPMC3545731.

35. Moretti R, Yen PS, Houe V, Lampazzi E, Desiderio A, Failloux AB, et al. Combining Wolbachia-induced sterility and virus protection to fight Aedes albopictus-borne viruses. PLoS Negl Trop Dis. 2018;12(7):e0006626. doi: 10.1371/journal.pntd.0006626. PubMed PMID: 30020933; PubMed Central PMCID: PMCPMC6066253.

36. Xi Z, Khoo CC, Dobson SL. Wolbachia establishment and invasion in an Aedes aegypti laboratory population. Science. 2005;310(5746):326-8.

37. Slatko BE, Luck AN, Dobson SL, Foster JM. Wolbachia endosymbionts and human disease control. Molecular and biochemical parasitology. 2014;195(2):88–95.

38. Xue L, Fang X, Hyman JM. Comparing the effectiveness of different strains of Wolbachia for controlling chikungunya, dengue fever, and zika. PLoS Neglected Tropical Diseases. 2018;12(7):e0006666.

39. Ant TH, Sinkins SP. A Wolbachia triple-strain infection generates self-incompatibility in Aedes albopictus and transmission instability in Aedes aegypti. Parasit Vectors. 2018;11(1):295. doi: 10.1186/s13071-018-2870-0. PubMed PMID: 29751814; PubMed Central PMCID: PMCPMC5948879.

40. Beebe NW, Pagendam D, Trewin BJ, Boomer A, Bradford M, Ford A, et al. Releasing incompatible males drives strong suppression across populations of wild and Wolbachia-carrying Aedes aegypti in Australia. Proc Natl Acad Sci U S A. 2021;118(41). doi: 10.1073/pnas.2106828118. PubMed PMID: 34607949; PubMed Central PMCID: PMCPMC8521666.

41. Ross PA, Robinson KL, Yang Q, Callahan AG, Schmidt TL, Axford JK, et al. A decade of stability for wMel Wolbachia in natural Aedes aegypti populations. PLoS Pathog. 2022;18(2):e1010256. doi: 10.1371/journal.ppat.1010256. PubMed PMID: 35196357; PubMed Central PMCID: PMCPMC8901071.

42. Handelsman J, Rondon MR, Brady SF, Clardy J, Goodman RM. Molecular biological access to the chemistry of unknown soil microbes: a new frontier for natural products. Chemistry & biology. 1998;5(10):R245–R9.

43. Cook JM, Butcher RD. The transmission and effects of Wolbachia bacteria in parasitoids. Researches on population ecology. 1999;41(1):15–28.

44. Salamon D, Zapała B, Krawczyk A, Potasiewicz A, Nikiforuk A, Stój A, et al. Comparison of iSeq and MiSeq as the two platforms for 16S rRNA sequencing in the study of the gut of rat microbiome. Applied microbiology and biotechnology. 2022;106(22):7671–81. Epub 2022/11/03. doi: 10.1007/s00253-022-12251-z. PubMed PMID: 36322250; PubMed Central PMCID: PMCPMC9628524.

45. Rueda LM. Pictorial keys for the identification of mosquitoes (Diptera: Culicidae) associated with dengue virus transmission. Walter Reed Army Inst Of Research Washington Dc Department Of Entomology, 2004.

46. Folmer O, Hoeh W, Black M, Vrijenhoek R. Conserved primers for PCR amplification of mitochondrial DNA from different invertebrate phyla. Molecular Marine Biology and Biotechnology. 1994;3(5):294–9.

47. Kim J, Hong J, Lim J-A, Heu S, Roh E. Improved multiplex PCR primers for rapid identification of coagulase-negative staphylococci. Archives of microbiology. 2018;200:73–83.

48. Caporaso JG, Kuczynski J, Stombaugh J, Bittinger K, Bushman FD, Costello EK, et al. QIIME allows analysis of high-throughput community sequencing data. Nature methods. 2010;7(5):335–6.

49. Yoon S-H, Ha S-M, Kwon S, Lim J, Kim Y, Seo H, et al. Introducing EzBioCloud: a taxonomically united database of 16S rRNA gene sequences and whole-genome assemblies. International journal of systematic and evolutionary microbiology. 2017;67(5):1613.

50. Edgar RC. Search and clustering orders of magnitude faster than BLAST. Bioinformatics. 2010;26(19):2460–1. doi: 10.1093/bioinformatics/btq461.

51. Mejia AJ, Jimenez L, Dutra HLC, Perera R, McGraw EA. Attempts to use breeding approaches in Aedes aegypti to create lines with distinct and stable relative Wolbachia densities. Heredity (Edinb). 2022;129(4):215–24. doi: 10.1038/s41437-022-00553-x. PubMed PMID: 35869302; PubMed Central PMCID: PMCPMC9519544.

52. Lubna, Rasheed SB, Zaidi F. Species diversity pattern of mosquitoes (Diptera: Culicidae) breeding in different permanent, temporary and natural container habitats of Peshawar, KP Pakistan. Braz J Biol. 2023;84:e271524. doi: 10.1590/1519-6984.271524. PubMed PMID: 37194758.

53. Jabeen A, Ansari J, Ikram A, Khan M, Tahir MA, Safdar M. A review of the geographical distribution of Aedes aegypti, Aedes albopictus and other Aedes species (Diptera: Culicidae) in Pakistan. Int J Mosq Res. 2019;6:90–5.

54. Ashfaq M, Hebert PD, Mirza JH, Khan AM, Zafar Y, Mirza MS. Analyzing mosquito (Diptera: Culicidae) diversity in Pakistan by DNA barcoding. PLoS One. 2014;9(5):e97268.

55. Calvez E, Guillaumot L, Millet L, Marie J, Bossin H, Rama V, et al. Genetic Diversity and Phylogeny of Aedes aegypti, the Main Arbovirus Vector in the Pacific. PLoS Negl Trop Dis. 2016;10(1):e0004374. Epub 2016/01/23. doi: 10.1371/journal.pntd.0004374. PubMed PMID: 26799213; PubMed Central PMCID: PMCPMC4723151.

56. Bel Mokhtar N, Maurady A, Britel MR, El Bouhssini M, Batargias C, Stathopoulou P, et al. Detection of Wolbachia infections in natural and laboratory populations of the Moroccan hessian fly, Mayetiola destructor (Say). Insects. 2020;11(6):340.

57. Juma EO, Allan BF, Kim C-H, Stone C, Dunlap C, Muturi EJ. Effect of life stage and pesticide exposure on the gut microbiota of Aedes albopictus and Culex pipiens L. Scientific reports. 2020;10(1):1–12.

58. Bennett KL, Gómez-Martínez C, Chin Y, Saltonstall K, McMillan WO, Rovira JR, et al. Dynamics and diversity of bacteria associated with the disease vectors Aedes aegypti and Aedes albopictus. Scientific reports. 2019;9(1):1–12.

59. Osei-Poku J, Han C, Mbogo CM, Jiggins FM. Identification of Wolbachia strains in mosquito disease vectors. PLoS One. 2012;7(11):e49922.

60. Zouache K, Raharimalala FN, Raquin V, Tran-Van V, Raveloson LHR, Ravelonandro P, et al. Bacterial diversity of field-caught mosquitoes, Aedes albopictus and Aedes aegypti, from different geographic regions of Madagascar. FEMS Microbiology Ecology. 2011;75(3):377–89.

61. Mancini M, Damiani C, Accoti A, Tallarita M, Nunzi E, Cappelli A, et al. Estimating bacteria diversity in different organs of nine species of mosquito by next generation sequencing. BMC microbiology. 2018;18(1):1–10.

62. Johnson M, Gómez A, Pinedo-Vasquez M. Land use and mosquito diversity in the Peruvian Amazon. Journal of Medical Entomology. 2008;45(6):1023–30.

63. Gonzalez-Ceron L, Santillan F, Rodriguez MH, Mendez D, Hernandez-Avila JE. Bacteria in midguts of field-collected Anopheles albimanus block Plasmodium vivax sporogonic development. Journal of medical entomology. 2003;40(3):371–4.

64. Campbell CL, Mummey DL, Schmidtmann ET, Wilson WC. Culture-independent analysis of midgut microbiota in the arbovirus vector Culicoides sonorensis (Diptera: Ceratopogonidae). Journal of Medical Entomology. 2004;41(3):340–8.

65. Takaya S, Kato Y, Katanami Y, Yamamoto K, Kutsuna S, Takeshita N, et al. Imported malaria at a referral hospital in Tokyo from 2005 to 2016: clinical experience and challenges in a non-endemic setting. The American Journal of Tropical Medicine and Hygiene. 2019;100(4):828.

66. Baldrian P, Kolařík M, Štursová M, Kopecký J, Valášková V, Větrovský T, et al. Active and total microbial communities in forest soil are largely different and highly stratified during decomposition. The ISME journal. 2012;6(2):248–58.

67. Zhang Q, Zhang Z, Lu T, Yu Y, Penuelas J, Zhu Y-G, et al. Gammaproteobacteria, a core taxon in the guts of soil fauna, are potential responders to environmental concentrations of soil pollutants. Microbiome. 2021;9(1):1–17.

68. Behar A, Yuval B, Jurkevitch E. Enterobacteria-mediated nitrogen fixation in natural populations of the fruit fly Ceratitis capitata. Molecular ecology. 2005;14(9):2637–43.

69. König H. Bacillus species in the intestine of termites and other soil invertebrates. Journal of applied microbiology. 2006;101(3):620–7.

70. Wu P, Sun P, Nie K, Zhu Y, Shi M, Xiao C, et al. A gut commensal bacterium promotes mosquito permissiveness to arboviruses. Cell host & microbe. 2019;25(1):101–12. e5.

71. Saab SA, Dohna Hz, Nilsson LK, Onorati P, Nakhleh J, Terenius O, et al. The environment and species affect gut bacteria composition in laboratory co-cultured Anopheles gambiae and Aedes albopictus mosquitoes. Scientific reports. 2020;10(1):1–13.

72. Sarwar MS, Jahan N, Ali A, Yousaf HK, Munzoor I. Establishment of Wolbachia infection in Aedes aegypti from Pakistan via embryonic microinjection and semi-field evaluation of general fitness of resultant mosquito population. Parasites & Vectors. 2022;15(1):1–13.

73. Gulraiz M, Alvi FM, Mustafa T, Razzaq A, Latif HS. Distribution of Aedes aegypti, Aedes albopictus and Culex sp. and detection of Wolbachia among them in city district Lahore. Journal of Fatima Jinnah Medical University. 2019;13(2):55–8.

74. Sinkins S, Braig H, Oneill SL. Wolbachia pipientis: bacterial density and unidirectional cytoplasmic incompatibility between infected populations of Aedes albopictus. Experimental parasitology. 1995;81(3):284–91.

75. Puerta-Guardo H, Contreras-Perera Y, Perez-Carrillo S, Che-Mendoza A, Ayora-Talavera G, Vazquez-Prokopec G, et al. Wolbachia in Native Populations of Aedes albopictus (Diptera: Culicidae) From Yucatan Peninsula, Mexico. Journal of Insect Science. 2020;20(5):16.

76. Shaikevich E, Bogacheva A, Rakova V, Ganushkina L, Ilinsky Y. Wolbachia symbionts in mosquitoes: Intra-and intersupergroup recombinations, horizontal transmission and evolution. Mol Phylogenet Evol. 2019;134:24–34. doi: 10.1016/j.ympev.2019.01.020. PubMed PMID: 30708172.

77. Ramírez-Puebla ST, Servín-Garcidueñas LE, Ormeño-Orrillo E, de León AV-P, Rosenblueth M, Delaye L, et al. Species in Wolbachia? Proposal for the designation of ‘Candidatus Wolbachia bourtzisii’,‘Candidatus Wolbachia onchocercicola’,‘Candidatus Wolbachia blaxteri’,‘Candidatus Wolbachia brugii’,‘Candidatus Wolbachia taylori’,‘Candidatus Wolbachia collembolicola’and ‘Candidatus Wolbachia multihospitum’for the different species within Wolbachia supergroups. Systematic and applied microbiology. 2015;38(6):390–9.

78. Ramirez-Puebla ST, Ormeno-Orrillo E, Vera-Ponce de Leon A, Lozano L, Sanchez-Flores A, Rosenblueth M, et al. Genomes of Candidatus Wolbachia bourtzisii wDacA and Candidatus Wolbachia pipientis wDacB from the Cochineal Insect Dactylopius coccus (Hemiptera: Dactylopiidae). G3 (Bethesda). 2016;6(10):3343-9. doi: 10.1534/g3.116.031237. PubMed PMID: 27543297; PubMed Central PMCID: PMCPMC5068953.

79. Sureshan SC, Mohideen HS, Nair TS. Gut Metagenomic Profiling of Gossypol Induced Oxycarenus laetus (Hemiptera: Lygaeidae) Reveals Gossypol Tolerating Bacterial Species. Indian Journal of Microbiology. 2022;62(1):54–60.

80. Moran NA, Degnan PH. Functional genomics of Buchnera and the ecology of aphid hosts. Molecular Ecology. 2006;15(5):1251–61.

81. Koch A. Endosymbiosis of Animals with Plant Micro-Organisms. Paul Buchner. The Quarterly Review of Biology. 1967;42(1):74–5. doi: 10.1086/405300.

